# The synergistic effects of the VH families and CH regions of multimeric IgM on its interaction with FcμR, and antigen

**DOI:** 10.1101/2022.05.19.492610

**Authors:** Wei-Li Ling, Samuel Ken-En Gan

## Abstract

As the primary response antibody with increasing interest as a therapeutic antibody format, IgM is also the largest antibody structure among the five major human isotypes. Spontaneously forming pentamer and hexamers, IgM has avidity effects that could compensate for weaker interactions, although steric hindrances can occur for certain epitopes. With recent evidence of the heavy chain constant region affecting antigen binding and the VH families of the V-regions affecting FcR engagement found on other isotypes, we investigated CDR-grafted Trastuzumab and Pertuzumab VH1-7 IgMs for biolayer interferometry. From our panel of the 14 IgM variants, the V-regions holistically affected FcμR binding, and the IgM C-region modulated Her2 engagements with contributions from the V-regions and influences from protein L binding at the Vκ. These findings revealed the oligomerization effect of IgMs to play a significant role in both FcμR and antigen binding that is distinct from the other isotypes that can guide the development and protein en-gineering of IgM therapeutics.

## Introduction

IgM is the primary response antibody isotype produced in the adaptive immunes response [1] and is structurally the largest multimeric antibody among the major human isotypes (IgM, IgA, IgD, IgG and IgE). IgM oligomerizes at its Cμ4 domain of the heavy chain constant (C)-region tailpiece to form pentamers in the presence of J-chains and hexamers without [2,3]. Through the oligomerization, IgM has avidity effects that can compensate for weaker antigen binding that may be present as monomeric antibody iso-types. An example of such compensation was observed between Pertuzumab and Trastuzumab IgG1s and IgMs [4] where while Trastuzumab IgG1 often showed stronger interactions with the Her2 antigen than Pertuzumab IgG1 [5], as IgMs, the steric clashes from binding multiple Her2 molecules due to epitope accessibility [6] resulted in Per-tuzumab IgM interacting with Her2 stronger than Trastuzumab IgM [7].

As the primary response antibody, IgM contributes to the mucosal immunity via the pIgR expressed on the epithelial cells that fa the transport of secretory IgM [8]. Yet, FcμR (also known as TOSO or FAIM3) that is expressed on lymphoid cells (B cells, T cells and NK cells) [9-11] to bind the IgM CH4 domain [12] is the main receptor for IgM to activate the signalling cascade via its immunoglobulin tail tyrosine (ITT) phosphorylation motif [13].

With recent evidence showing that the antibody regions can cross-influence one an-other such as in the variable (V-) regions, particularly the framework regions (FWR), can influence the FcR binding sites (and thus FcR binding) at the heavy chain constant regions (CH) for IgA [14,15], IgE[16], and IgG1 [17], a holistic [18] investigation for multimeric IgMs can elucidate the contribution of oligomerization. Considering how the non-antigen interacting regions such as the signal peptide [19] impact antibody production (apart from the obvious VH-VL pairing on production [20]) and how the different regions such as within the V- and C-regions can come together to form amino acid stretches and pockets that can bind non-antigens [21,22], there remains much to investigate at the whole anti-body level and the effects of such oligomerization.

While most of the current antibody therapeutics are IgGs, the interest in the other isotypes such as IgM has increased in recent years. The contributory role of IgM in viral immunity was found in the recent SARS-CoV-2 pandemic [23-26] while the promise of IgM as a therapeutic antibody was shown through the listed company IGM Biosciences currently with therapeutic IgM candidates undergoing phases 1 and 2 clinical trials (ClinicalTrials.gov Identifier: NCT04553692 & CT04082936). It is important to understand in detail how the different regions in IgM interact and contribute to the antibody functions, considering the added uniqueness of IgM’s oligomerization.

With the further complication of antibody superantigens that bind to antibody re-gions [27,28] to modulate antigen and receptor engagement [28], there is a further need to determine how multimeric IgM can be influenced by superantigen binding (particularly Protein L or PpL which binds the Vκ light chain).

To investigate these effects, a panel of 14 variable heavy (VH) family variants of Trastuzumab and Pertuzumab (where the complementarity determining regions (CDRs) of the two clinical antibodies were grafted onto the VH1-7 family FWRs [17,19]) and their binding to Her2 antigen and FcμR receptor measured using Bio-Layer Interferometry (BLI) measurements.

## Result

### 2.1. BLI measurements of FcμR to Pertuzumab and Trastuzumab IgM variants

To measure the distal allosteric effects of the V-region VH family FWRs on the FcμR binding site at the IgM CH, the grafted Trastuzumab and Pertuzumab CDRs to the seven VH FWR were recombinantly connected to the IgM C-region for expression. All the VH1-7 variants of both Trastuzumab and Pertuzumab were produced successfully, albeit at different concentrations. Given that no J-chain was provided, the expected spontaneous formation of predominantly hexamers was estimated from the size exclusion chromatography (SEC) and electron microscopy (EM) on selected samples from our panel shown in Supplementary Figures S1-3.

With the expected multivalency of IgM, the 2:1 fitting models of the biolayer interferometry (BLI) Octet Red® software were utilized over the 1:1 fitting model. From the results where antibodies of the same VH family showed contrasting binding kinetics, additional factors beyond the VH FWR were shown to influence the interaction between the IgMs and FcμR. The results showed that the FWRs and CDRs of both the heavy and light chains synergistically affected the binding site at the CH.

Among the Pertuzumab VH1-7 variants (Figure 1), PVH1-IgM was the only Pertuzumab variant with significant interactions with FcμR, yielding KD 1 and KD 2 values of 45 and 29 nM, respectively. For the Trastuzumab variants, four out of the seven VH families bound FcμR with KD values ranging from 35 to 25740 nM. The strongest to weakest interacting Trastuzumab IgMs were in the order of VH7, 6, 5 & 2. As all the VH variants within the respective antibody models had the same Trastuzumab or Pertuzumab Vκ1 light chain, the light chain was ruled out as a single contributing partner to FcμR engagement.

**Figure 1.**
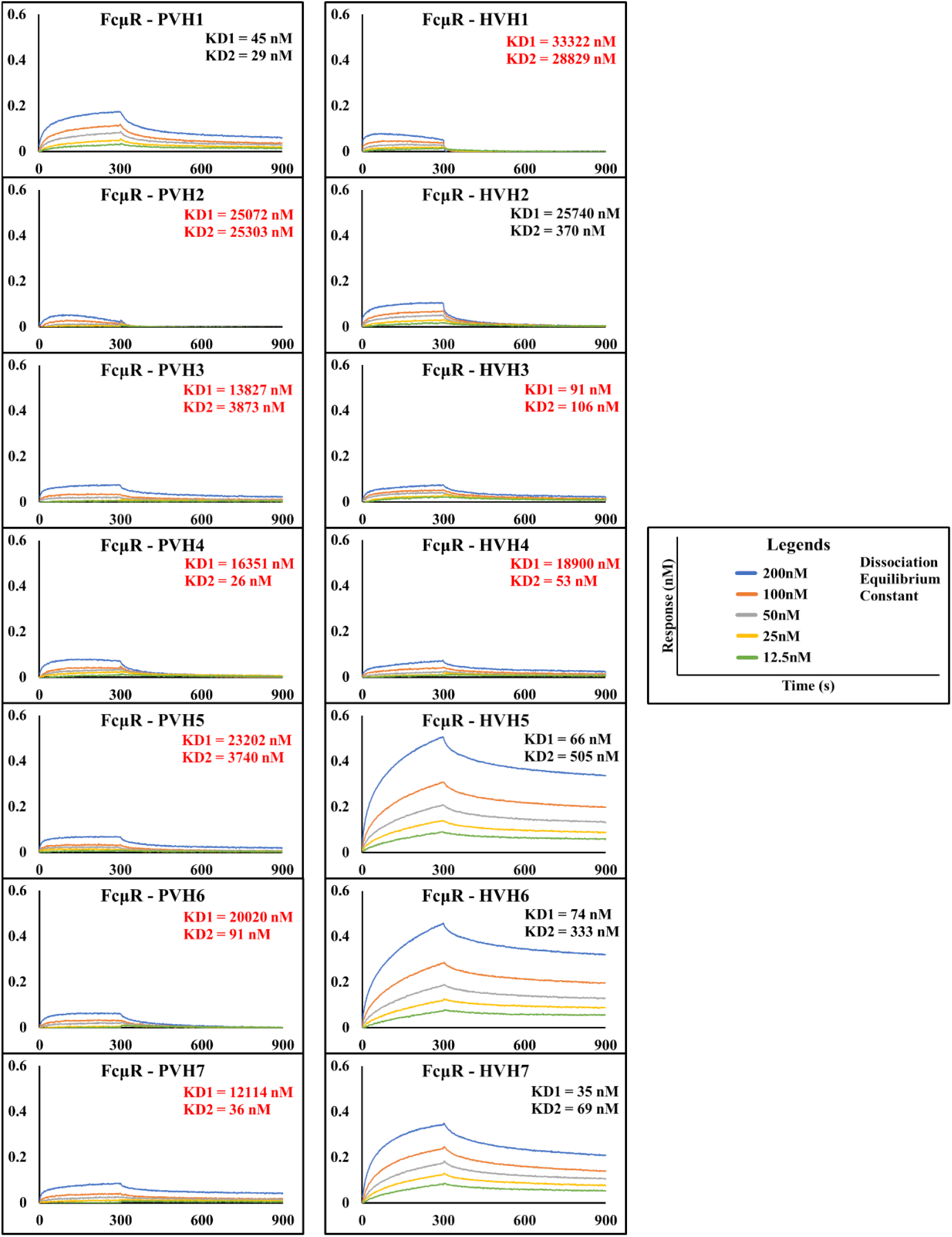
BLI measurements of the different concentrations of Pertuzumab (PVH1-7) and Trastuzumab (HVH1-7) IgM variants to FcμR bound to the AHC biosensor. Two KD values are shown according to the 2:1 fitting model. The KD values highlighted in red were deemed unreliable due to low R2 values (<0.9) and/or had a poor response (<0.1). KD values were rounded off to the nearest whole number.

### 2.2. BIL measurements of Pertuzumab and Trastuzumab IgM variants to Her2

As a control for protein folding and IgM functionality, measurements with the known Pertuzumab and Trastuzumab antigen – Her2 – were performed using two im-mobilization methods within the Octet Red96® system. The first method used Protein L (PpL) biosensor to capture the Vk1 light chains of the IgMs (Figure 2) at the FWR1 [21], while the second method used biotinylated anti-IgM antibodies (recognizing IgM CH) immobilized with Streptavidin (SA) biosensors (Figure 3). Both immobilization methods would yield results that took into consideration the avidity effects of IgM, the need for cor-rect folding of the IgM CH for recognition by anti-IgM, and the possible modulating effects of PpL on Her2 antigen binding. Although the Pertuzumab IgM VH variants (except for Pertuzumab VH1 IgM) did not interact with FcμR (Figure 2), all the Pertuzumab IgM variants were successfully captured by the anti-IgM antibodies and further interacted with Her2 showing a relatively narrow and expected nM range of KD1 and KD2 values (Figure 3). The measurements of the an-ti-IgM captured method were more consistent than those of PpL captured Pertuzumab IgMs to Her2 (Figure 2) showing modulating effects from the PpL engagement.

**Figure 2.**
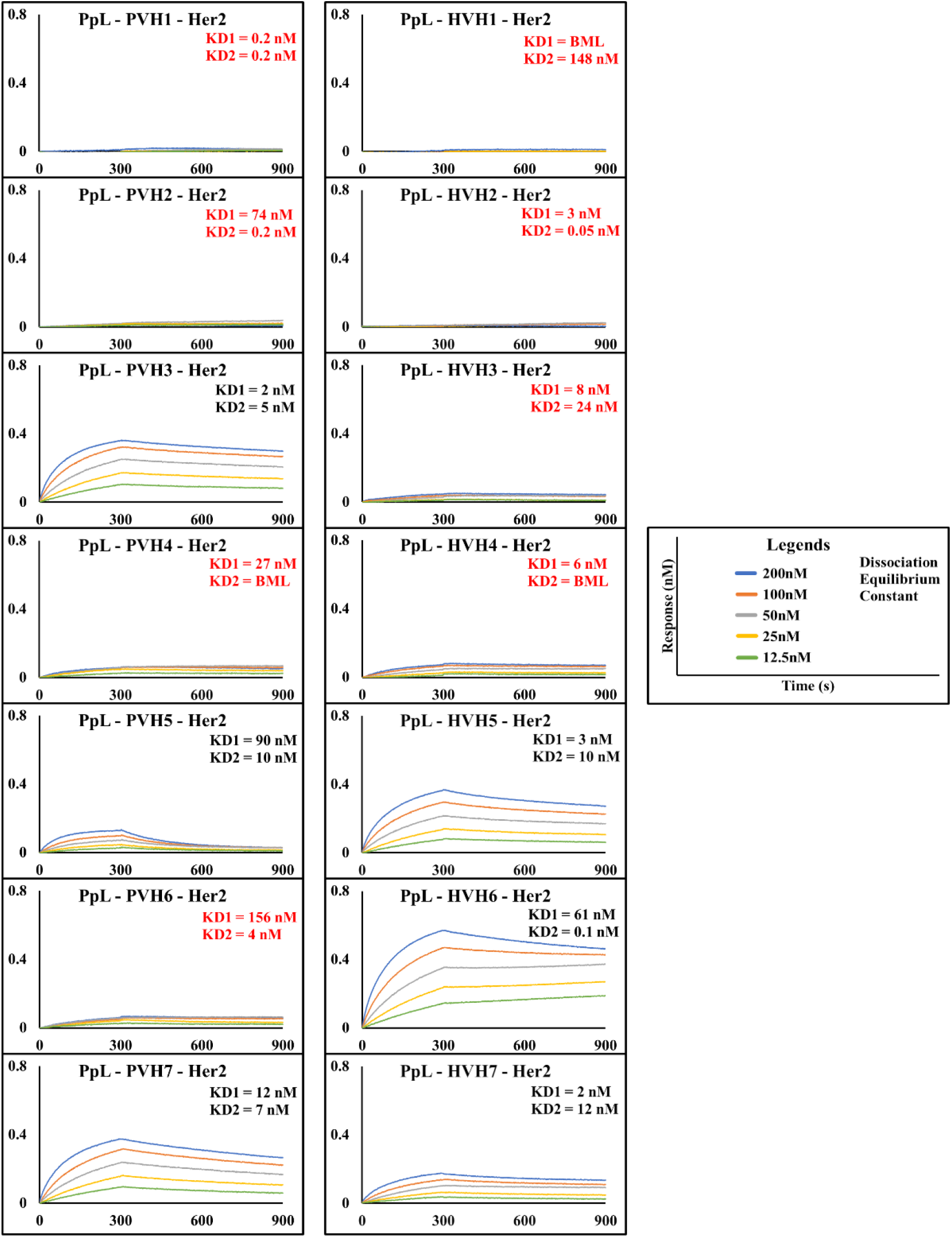
BLI measurements of the different concentrations of Pertuzumab (PVH1-7) and Trastuzumab (HVH1-7) IgM variants to Her2 using the Protein L (PpL) capture method. Two KD values are shown according to the 2:1 fitting model. The KD values highlighted in red were considered unreliable due to low R2 values (<0.9) and/or poor response (<0.1). KD values were rounded off to the nearest whole number. BML stands for “beyond ma-chine/measurement limit” where KD values generated are <1.0 × 10-12. The loading graph of each variant is shown in Supplementary Figures S4 and S5).

**Figure 3.**
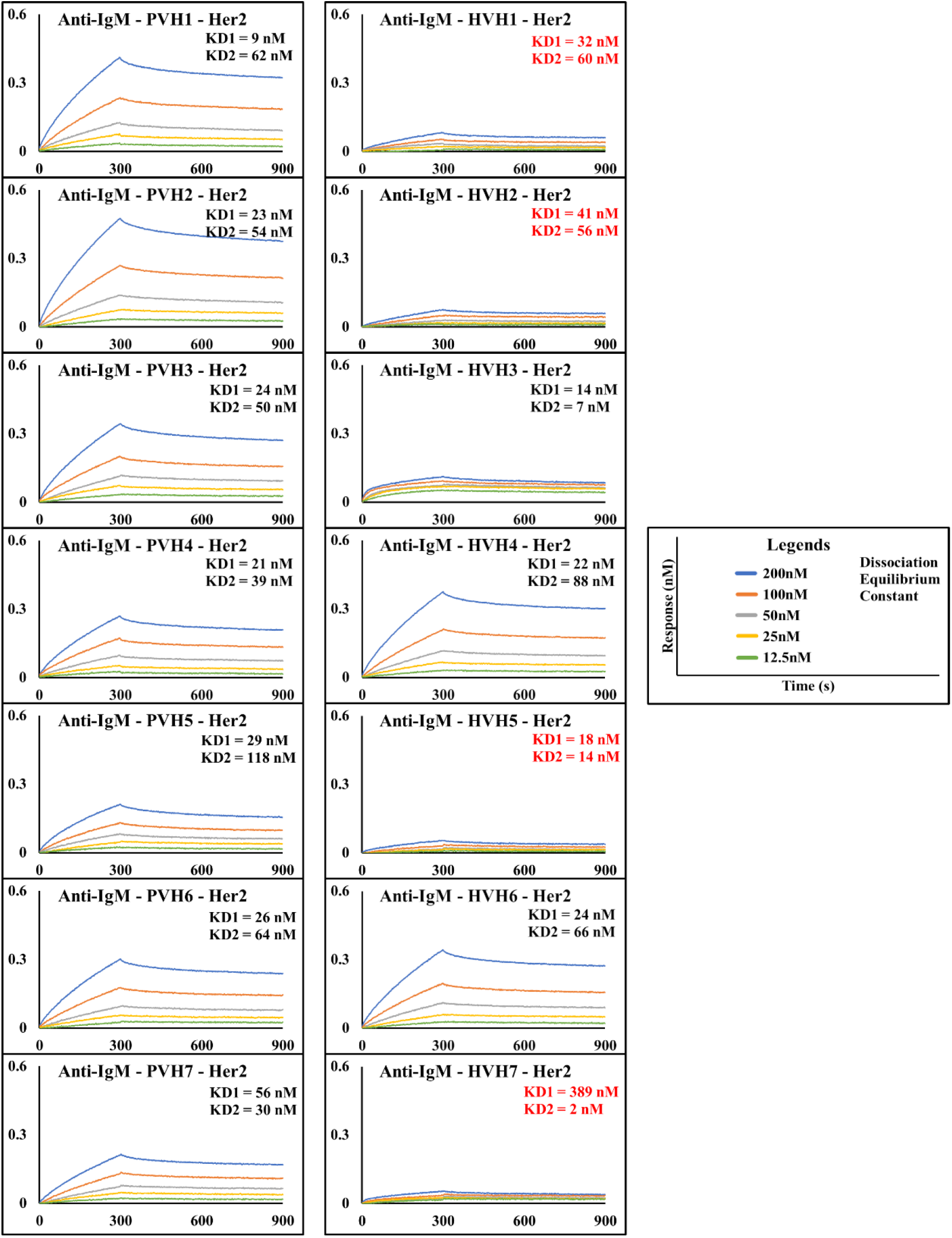
BLI measurements of the different concentrations of Pertuzumab (PVH1-7) and Trastuzumab (HVH1-7) IgM variants to Her2 using the anti-IgM capture method. Two KD values are shown according to the 2:1 fitting model. The KD values highlighted in red were considered unreliable due to low R2 values (<0.9) and/or low response (<0.1). KD values were rounded off to the nearest whole number. The loading graph of each variant is shown in Supplementary Figures S6 and S7).

Using the PpL immobilization method shown in Figure 2, three out of the seven VH Pertuzumab IgM variants interacted with Her2 with KD values ranging from 2 to 90 nM with the strongest to weakest interacting IgMs as VH3, 7 & 5.

Of the Trastuzumab IgM variants, three out of the seven VH families interacted with the Her2 with lower KD values (stronger interactions) ranging from 0.1 to 61 nM. Com-pared to the Pertuzumab IgMs with the order of VH3,7 and 5 for strongest to weakest bind-ers, the Trastuzumab IgMs were in the order of VH5/7 & VH6 (based on the average KD1 and KD2 values). VH5 and VH7 FWRs from both Pertuzumab and Trastuzumab IgM var-iants consistently interacted with Her2, with the former showing higher variability.

Anti-IgM captured IgM showed that all Pertuzumab VH IgM variants bound to Her2 with KD values ranging from 9 to 118 nM with VH4, VH1/2/3, VH6/7 & VH5 in the order of the strongest to the weakest binders (Figure 3). Among the seven Trastuzumab IgM vari-ants, only the VH3, 6, and 4 (in order of lowest KD to highest) yielded reliable KD1 and 2 averages ranging from 7 to 88 nM. It is noteworthy that the VH6 FWR on both Trastuzumab (HVH6) and Pertuzumab (PVH6) showed relatively consistent KD1 and KD2 values.

## Discussion

Through the CDR grafting of two well-studied cancer therapeutic antibodies, Pertuzumab and Trastuzumab, the VH FWRs previously affected CH-FcR interaction on oth-er human antibody isotypes [4,14-17,19,22]. Here, we set out to investigate the possible ef-fects of the seven VH families on the distal binding of IgM CH to its receptor FcμR by the recombinant expression of Pertuzumab and Trastuzumab IgM VH variants. These whole IgMs were made by stitching the previous VH variants onto the human IgM CH [4] that spontaneously formed multimers consistent with our previous work on IgM [3,29,30] and from our SEC and EM analyses of selected samples (Supplementary Material Figure S1-3).

Since the binding to its respective FcR by the antibody affects its activation of downstream signalling on the immune cells, impaired receptor FcμR engagement can lead to lower efficacy in IgM therapeutics. Found on lymphocytes such as natural killer cells, B cells, and T cells [9-11], FcμR binds IgM at the CH. In agreement with recent investigations that showed the V-regions (Both VH and VL in combination) influencing FcR binding sites on the CH region for IgA [14] [15], IgE [16], IgG1 [17], similar effects were found here for IgM. With the unique oligomerization of IgM from the other human isotypes in oligomer-ization, the allosteric communications between the V- and C-regions were found ampli-fied to modulate IgM- FcμR interactions with all the Pertuzumab IgM except for VH1 and only three Trastuzumab IgM variants having significant interactions with FcμR (Figure 1).

The IgM variants in our panel interacted with FcμR within the expected ∼10 nM avidity interaction [31] from a previous study that utilized a different measurement method with the assumption of 1:1 stoichiometry [31]. Our attempts with the Octet Red® 1:1 fitting model could not yield reasonable KD values and fitting curves, requiring the 2:1 fitting model to yield KD values that were ∼10 nM.

From the lack of patterns within VH-FWR families on FcμR-CH engagement, there were clear contributions from the CDRs of both VH and Vκ given that the same Vκ1 FWR was used for both Pertuzumab and Trastuzumab. Such combinatorial V-region effects were shown where the VH1 FWR, producing a FcμR binding IgM with Pertuzumab CDRs (PVH1) and light chain, did not show significant FcμR engagement with the Trastuzumab CDRs (HVH1) and light chain. The reversal of such patterns was also observed with VH7, 5, and 6 FWRs of Trastuzumab but not Pertuzumab interacting with FcμR.

As a control for correct folding and functionality of the recombinant IgMs were eval-uated using PpL (Vκ capture) and anti-IgM (IgM CH regions capture) immobilization methods. Through the successful loading of the IgMs by the two capture methods (Sup-plementary Figure S4-7) demonstrating recognizable IgM binding sites, the lack of interac-tions FcμR exhibited by the majority of our panel of IgMs was not due to misfolded or non-functional recombinant IgMs.

The two immobilization capture methods also allowed investigations into the proximal modulation of superantigen PpL binding the Vκ FWRs on antigen Her2 binding. Although both Trastuzumab and Pertuzumab bind Her2 as their antigen, they bound at different epitopes that were previously shown to affect accessibility and cause steric hin-drances [6,7] for the multivalency of IgM. Further, it is reasonable to expect that the bind-ing of superantigens [14,32], particularly at the V-regions could therefore modulate anti-gen recognition due to the close packing of the V-regions for IgM.

From the PpL capture method results (Figure 2), only PVH3, 5 & 7 of the Pertuzumab IgM variants interacted with Her2 whereas all the Pertuzumab IgM variants could with the anti-IgM capture method (Figure 3). With the PpL immunomodulating the effects, the importance of the microbial superantigens from normal flora (discussed in [32] and [27]) is also expected to extend to other isotypes given the binding at the Vκ-FWR1 [21] that is utilized for both Trastuzumab and Pertuzumab.

The log-fold difference between Pertuzumab VH5 IgM and Trastuzumab VH5 IgM within the anti-IgM immobilization results were in the same trend as the differences ob-served previously between Pertuzumab and Trastuzumab IgG1s [5]. Compared to the VH5 variants, the compensatory effect of Pertuzumab VH7 IgM having similar KD values with that of Trastuzumab VH7 IgM reflected the avidity effects [4] and even epitope acces-sibility issues to bind Her2 as discussed previously [6,7].

Analysing the Trastuzumab dataset, not all the Trastuzumab VH IgMs interacted with Her2 regardless of the immobilization method. Clear interferences from PpL in pre-venting the HVH1-4 from binding Her2 were also observed. Curiously, HVH1 and 2 were consistently unable to bind Her2 despite the capture method, but the same CDR-grafting on Pertuzumab IgM captured by anti-IgM could. Given that the two HVH V-regions could bind to Her2 when made into IgG1s [17], IgAs [14], and IgEs [16] using the same V-regions and light chains, there was a clear structural interference from the IgM CH oligomeriza-tion on the IgM-Her2 binding. With the pattern for the rest of the Trastuzumab VH vari-ants binding Her2 when captured by PpL or anti-IgM, the recombinant IgMs were cer-tainly functional, and the modulation was amplified by the oligomerization of IgM CH.

With the growing importance of IgM as a therapeutic antibody and its use in diag-nostics (such as in haemagglutination kits), uncovering the mechanisms of these factors and how they can be mitigated by the choice of FWR in CDR-grafting in humanization [33] would be crucial for the efficacy of IgM for therapeutic and diagnostic uses.

Our panel showed the VH7 FWR to be a potentially suitable FWR for CDR grafting for retaining the bulk of the IgM antigen binding abilities although this consensus was absent for FcμR engagement. Certainly, the oligomerization of IgM amplified the distal re-gional effects between the V- and C-regions, suggesting the need for future studies to in-vestigate IgM candidates individually yet holistically due to the combinatorial factors in-volved in mechanisms that are yet to be fully elucidated. Such investigations would also need to include the proximal effects elicited by PpL on Her2 binding as found for IgEs and IgAs of the same V-regions [14,16] suggesting unique characteristics of IgM CH oligomer-ization. Given that IgM is the primary response antibody, our findings here lend support to the hypothesis for class-switching as a mechanism for affinity maturation that may be taken into consideration during the humanization during antibody engineering [33].

In conclusion, we found that not all IgMs would engage FcμR efficiently to elicit the necessary downstream immune effects and that the oligomerization of the IgM could module antigen-binding by V-regions. With further contributory effects from V-region su-perantigens such as PpL to affect antigen binding, the effects were amplified in multimeric IgM. Our findings here also support the use of CH engineering in antibody engineering and the hypothesis that class switching may be a mechanism for affinity maturation in humans.

## Materials and Methods

The recombinant IgM variants with Trastuzumab (PDB 1N8Z) and Pertuzumab (PDB 1S78) sequences were generated by subcloning restriction enzyme (EcoRI and NheI) digested gene from the IgG1 VH1-7 variants [17,19] to the IgM CH [4] and signal peptides [19,21,34,35]. The light chains were created by grafting the respective Pertuzumab and Trastuzumab Vκ1 genes with the κ-constant region. Briefly, the plasmids (one heavy chain and one light chain) were transformed into DH5α competent cells [37] separately for plasmid production and purified using the column-based plasmid extraction methodol-ogy [36].

The heavy and light chain plasmids were then co-transfected into HEK293 cells us-ing PEI MAX® (Cat: 24765-1, PolyScience). The production supernatant was harvested af-ter 14 days post-transfection and purified using AKTA equipped with a PpL affinity col-umn (Cat: 17547815, Cytiva). as previously optimized [20] and [38]. The purified recom-binant IgMs were then subjected to HiLoad Superdex 200 pg preparative SEC columns (Cat: 28989335, Cytiva) for the collection of the spontaneously formed hexamer fractions as previously described work [4] where only the multimeric fraction was collected for analysis.

### BLI Measurements

KD measurements of Fc-tagged Human FcμR (Cat: 13556-H02H, SinoBiological) immobilised on Anti-Human IgG Fc (AHC) (Cat: 18-5060, Sartorius) biosensors were performed on the Octet Red96® system with the loading threshold set at 1.0 nm and utilizing the recombinant Pertuzumab and Trastuzumab IgM VH whole antibodies variants in five serial diluted concentrations (12.5 to 200nM) solutions.

KD measurements of the Pertuzumab and Trastuzumab IgMs to their Her2 antigen (Cat: 10004-HCCH, SinoBiological) were performed by first immobilizing them on Protein L (Cat: 18-5085, Sartorius) and CaptureSelect™ Biotin Anti-IgM Conjugate (Cat: 7102892500, Thermofisher Scientific) bound Streptavidin (SA) biosensors (Cat: 18-5019, Sartorius) (loading threshold set at 1.0 and 0.5 nm, for Protein L and Anti-IgM SA biosen-sor, respectively) and subjected to the five serially diluted Her2 solutions (12.5 to 200nM).

The program and steps used were described as followed. The Octet Acquisition v10.0 program was used to set the following steps for KD measurements: Pre-conditioning (0.2 M glycine, pH 1.52 & 10x kinetic buffer (KB) (Cat 18-1105, Sartorius), 30 s); Initial Baseline (10x KB, 60 s); Loading (ligands, 600s with threshold) Baseline (10x KB, 120 s); Association (analyte, 300 s); Dissociation (10x KB, 600 s); Regeneration (0.2 M glycine, Ph 1.52 & 10x KB), 30 s). The equilibrium dissociation constant (KD) values were automatically calculated from the ka and kd using the Data Analysis v10.0 program using the 2:1 fitting model (The 1:1 model did not yield reasonable KD values). KD measurements were considered reliable if the R2 value is above 0.9 and the response value is above 0.1. All measurements were performed in triplicates as previously performed [4,5,14-19,21,22,28]. The 2:1 fitting models were used to analyse the IgM interactions and are shown in Results as well as Supplementary Materials.

## Acknowledgements

This work was supported by the Joint Council Office, Agency for Science, Technology, and Research, Singapore under Grant number JCO1334i00050 and by APD SKEG Pte Ltd. We would like to acknowledge the A*STAR Microscopy platform for the EM and cryo-EM analysis of IgM structures.

## Author Contributions

**Conceptualization, Methodology, Investigation, Writing was by W.L.L. and S.K-E.G. Funding Acquisition, S.K-E.G. Supervision, S.K-E.G**.

## Data availability

The datasets GENERATED/ANALYZED for this study are available upon request.

