## Supplementary material for "The synergistic effects of the VH families and CH regions of multimeric IgM on its interaction with FcμR, and antigen": Supp

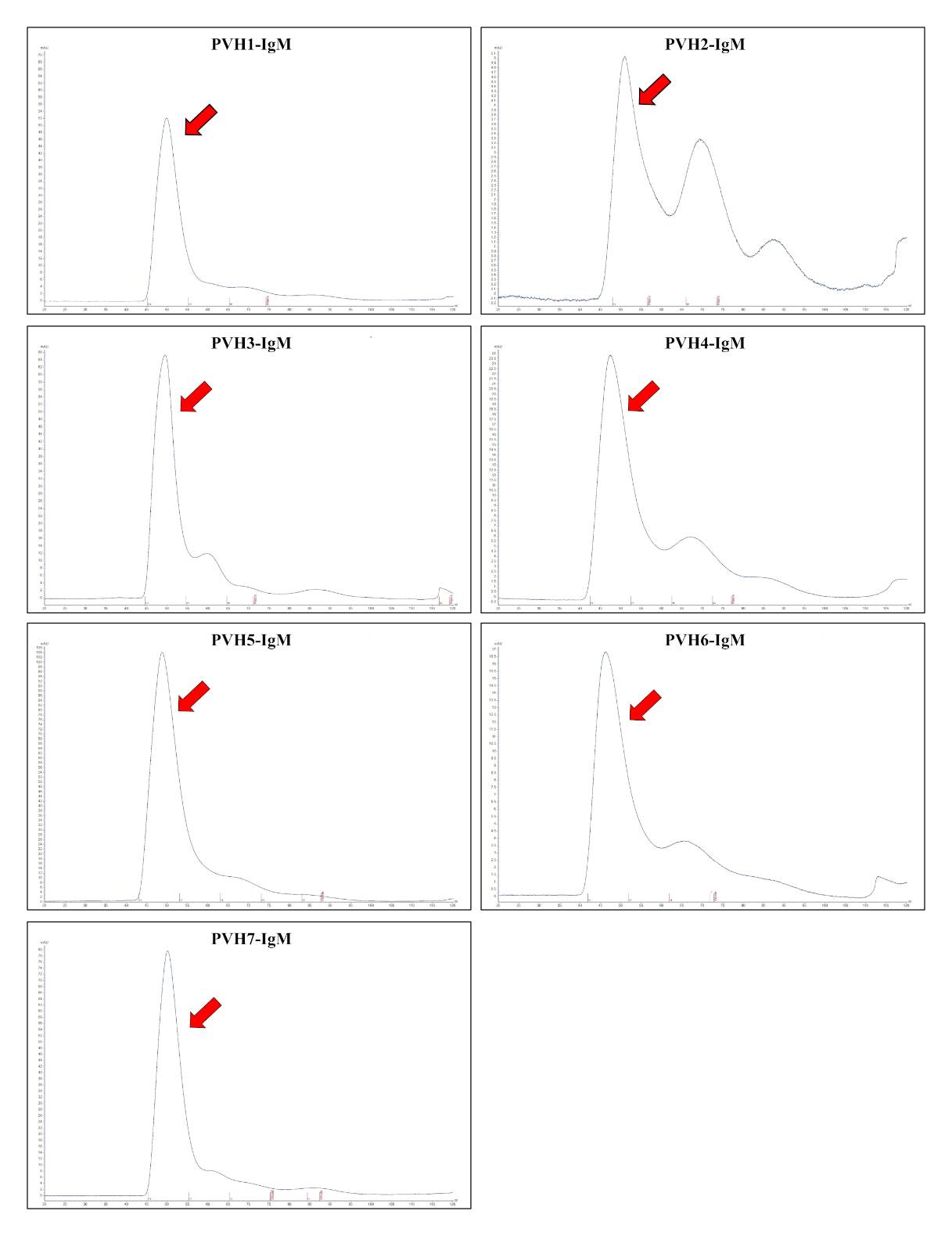

**Supplementary Figure S1. Size exclusion chromatogram of PVH1-7 IgM variants. The collection of the multimeric fraction were as indicted the by the red arrow.**

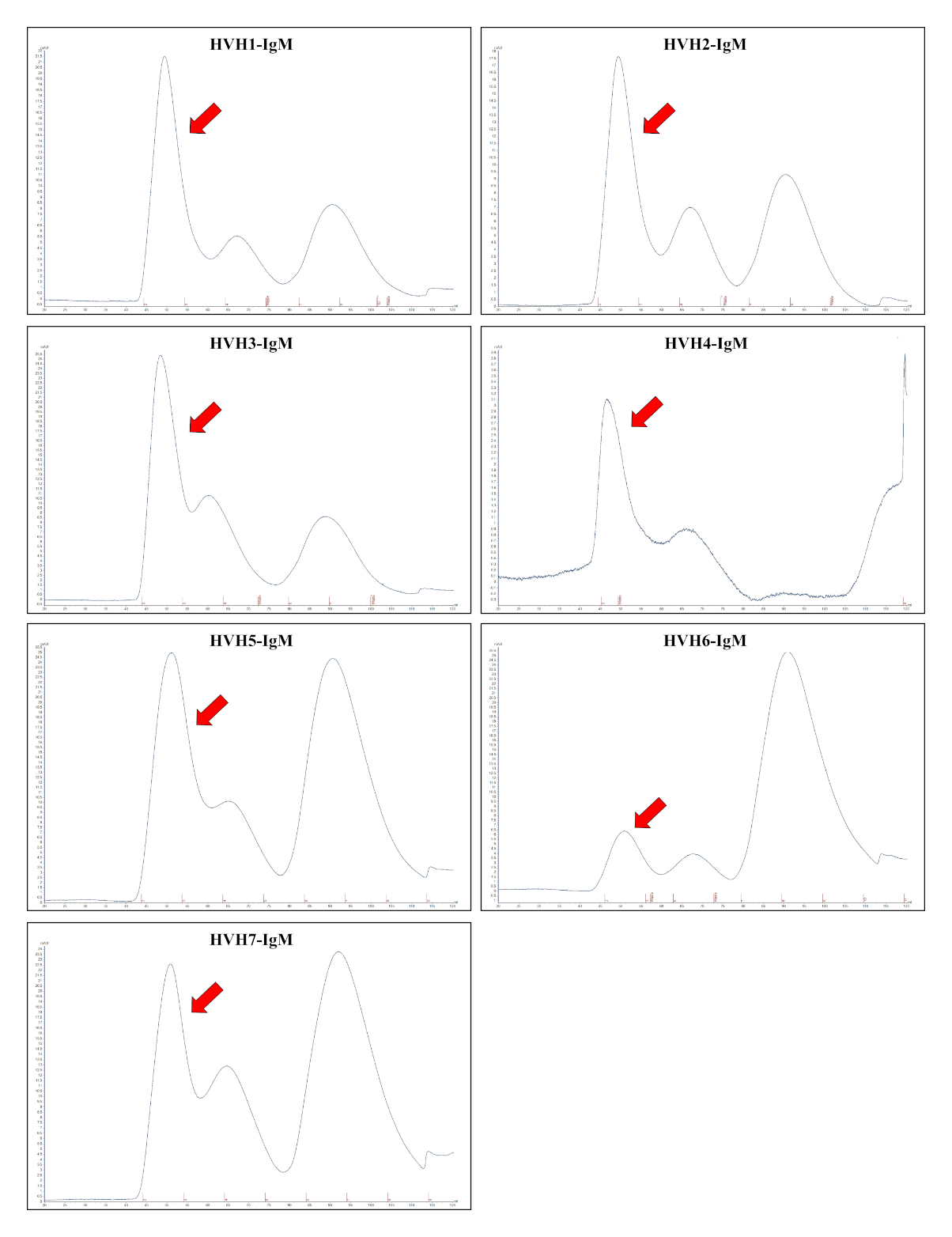

**Supplementary Figure S2. Size exclusion chromatogram of HVH1-7 IgM variants. The collection of the multimeric fraction were as indicted the by the red arrow.**

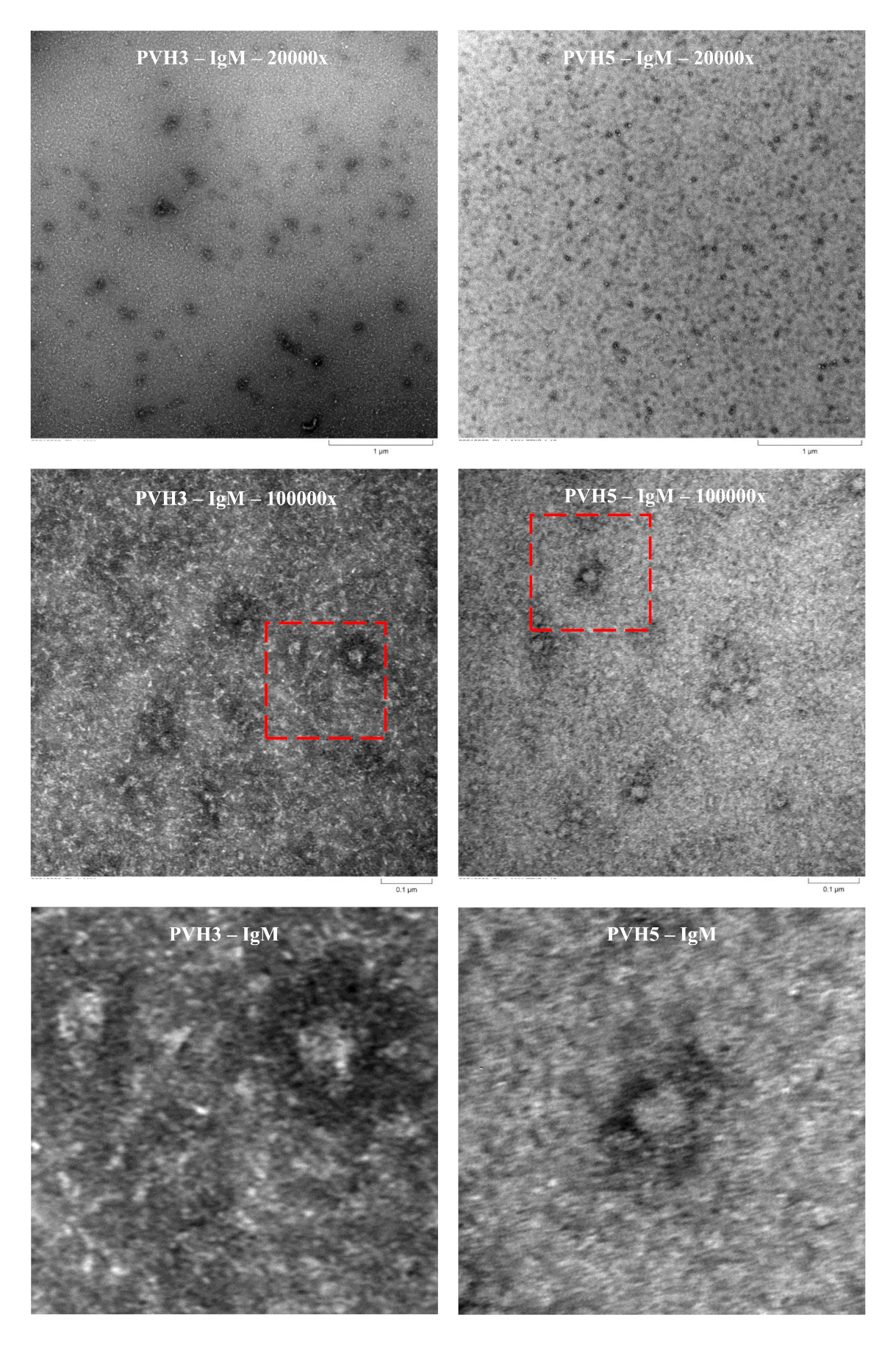

**Supplementary Figure S3. Samples, PVH1 and PVH3 IgM have been negatively stained with Uranyl Acetate and images were acquired using 80kV TEM (RT). The scale bars are indicated, except for the two images at the bottom that are 4x enlarged zoom of the 100000x images above.**

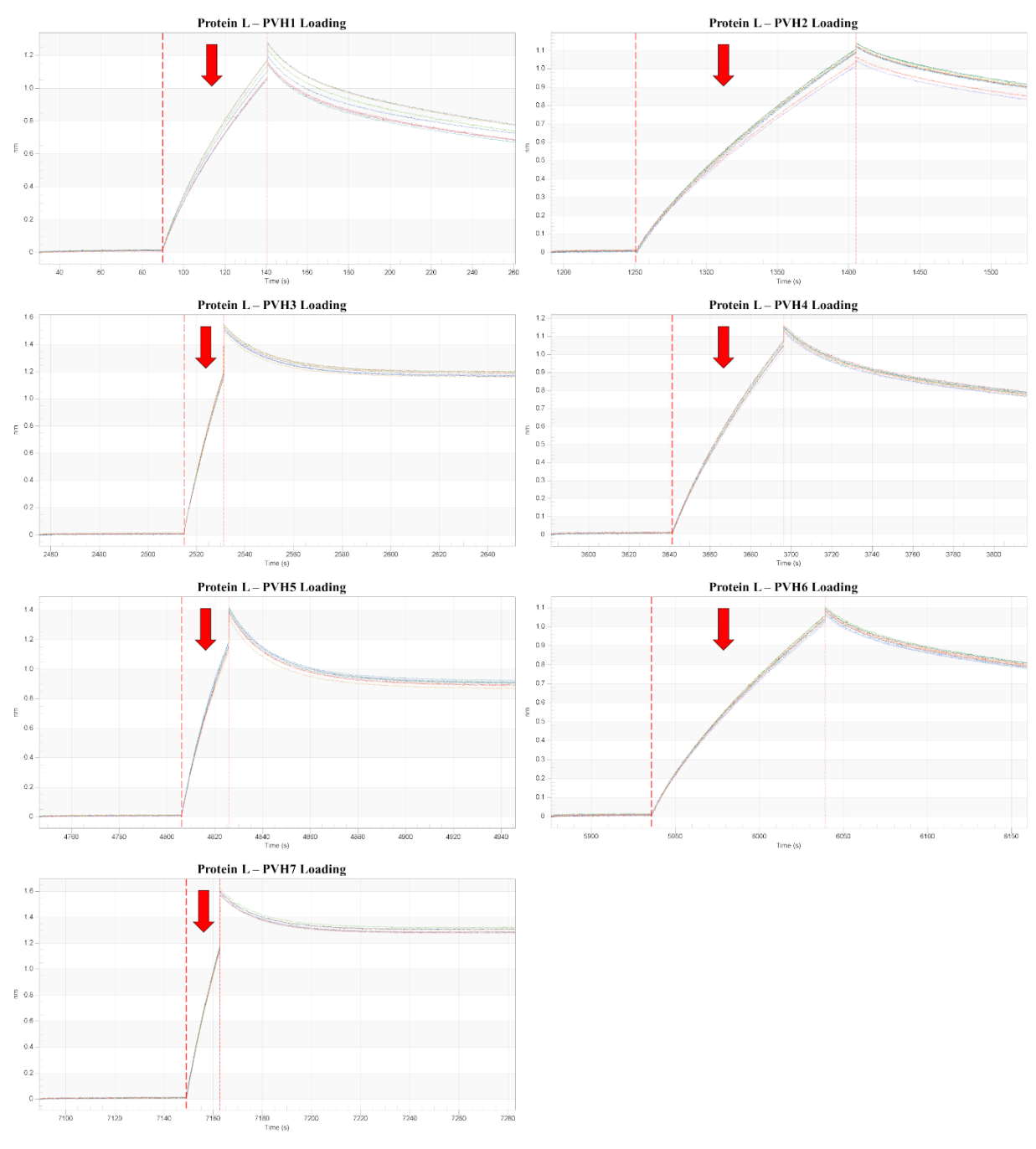

**Supplementary Figure S4. Loading graph of PVH1-7 on Protein L biosensor.**

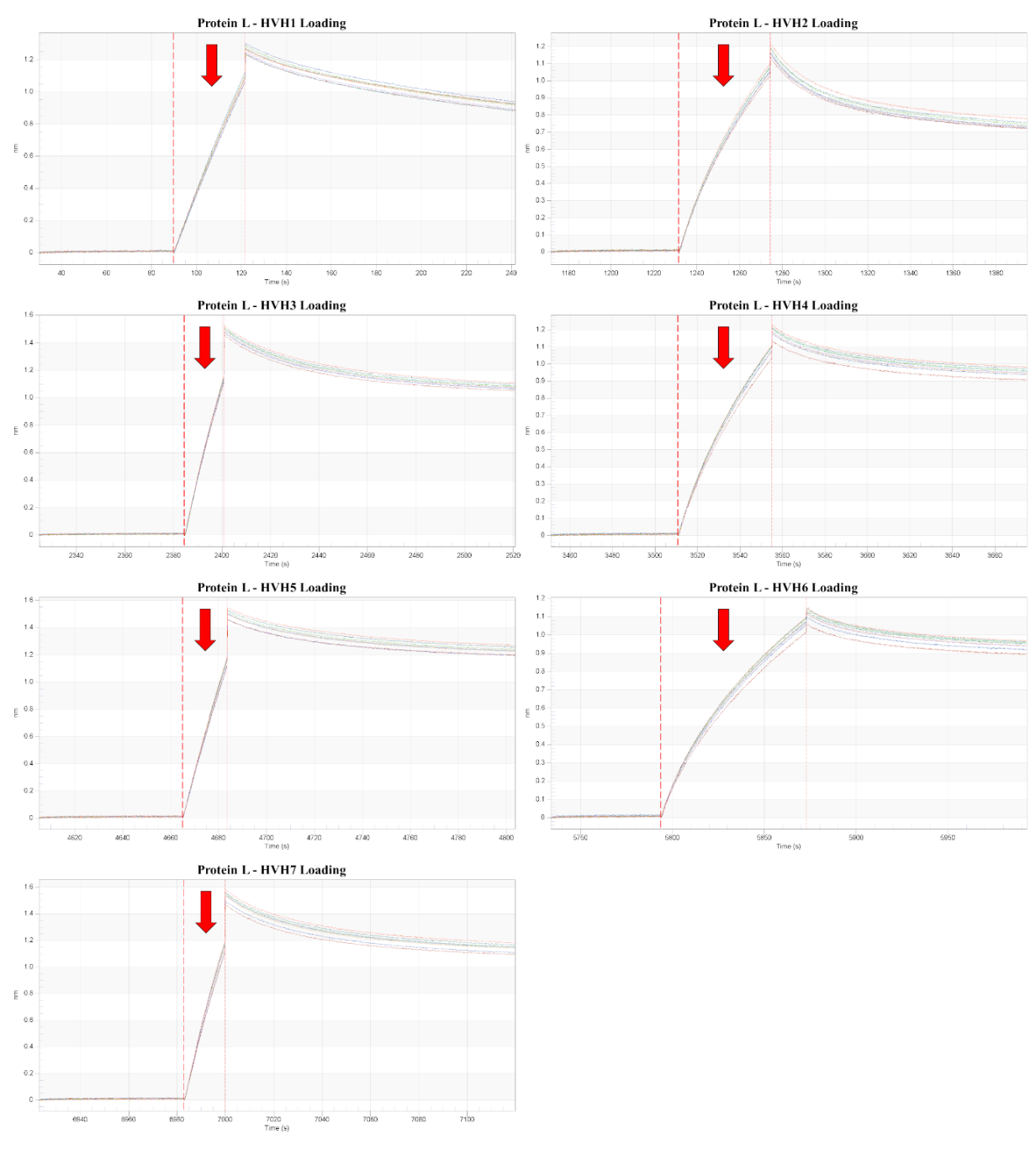

**Supplementary Figure S5. Loading graph of HVH1-7 on Protein L biosensor.**

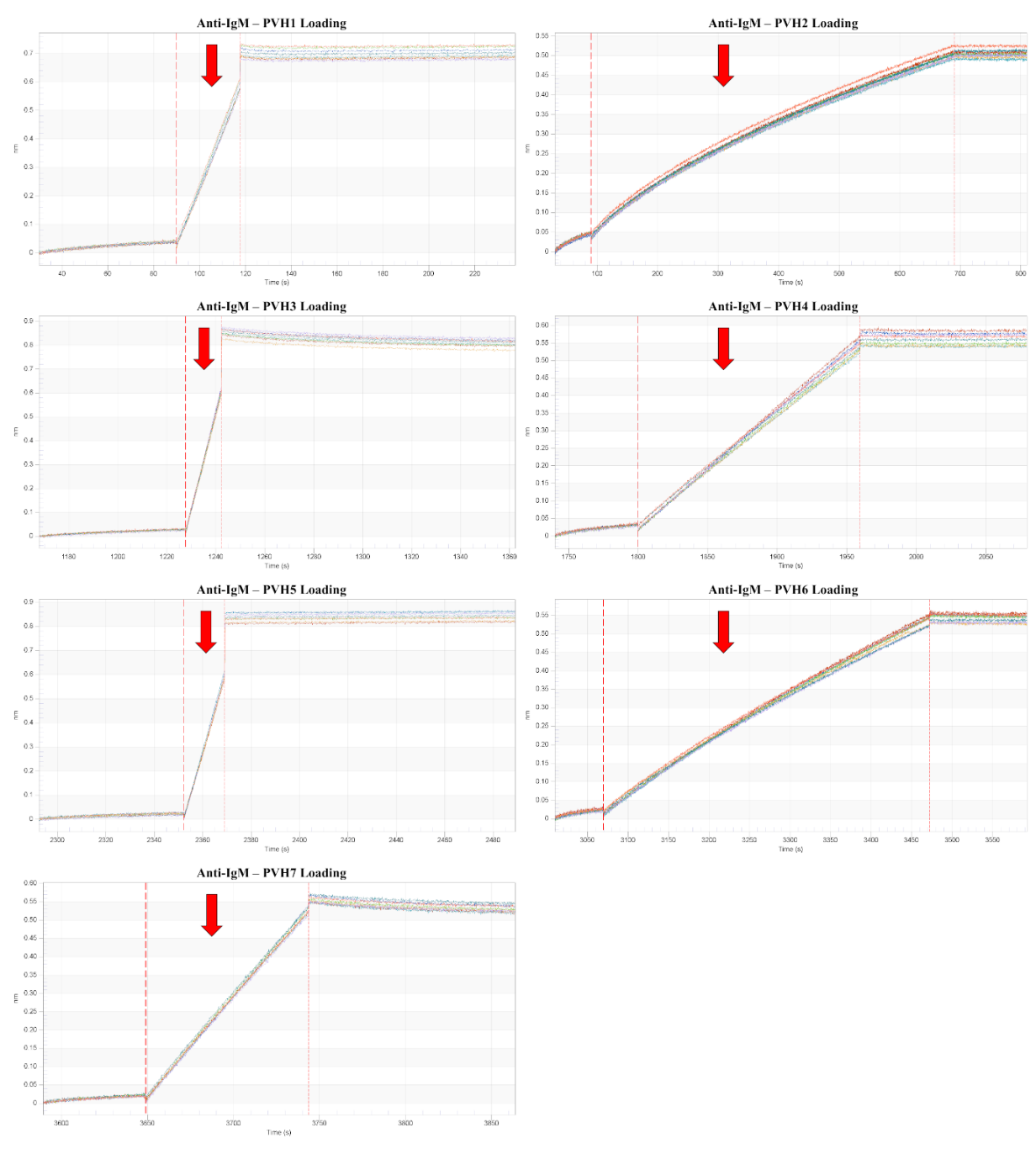

**Supplementary Figure S6. Loading graph of PVH1-7 on Anti-IgM bound on SA biosensor.**

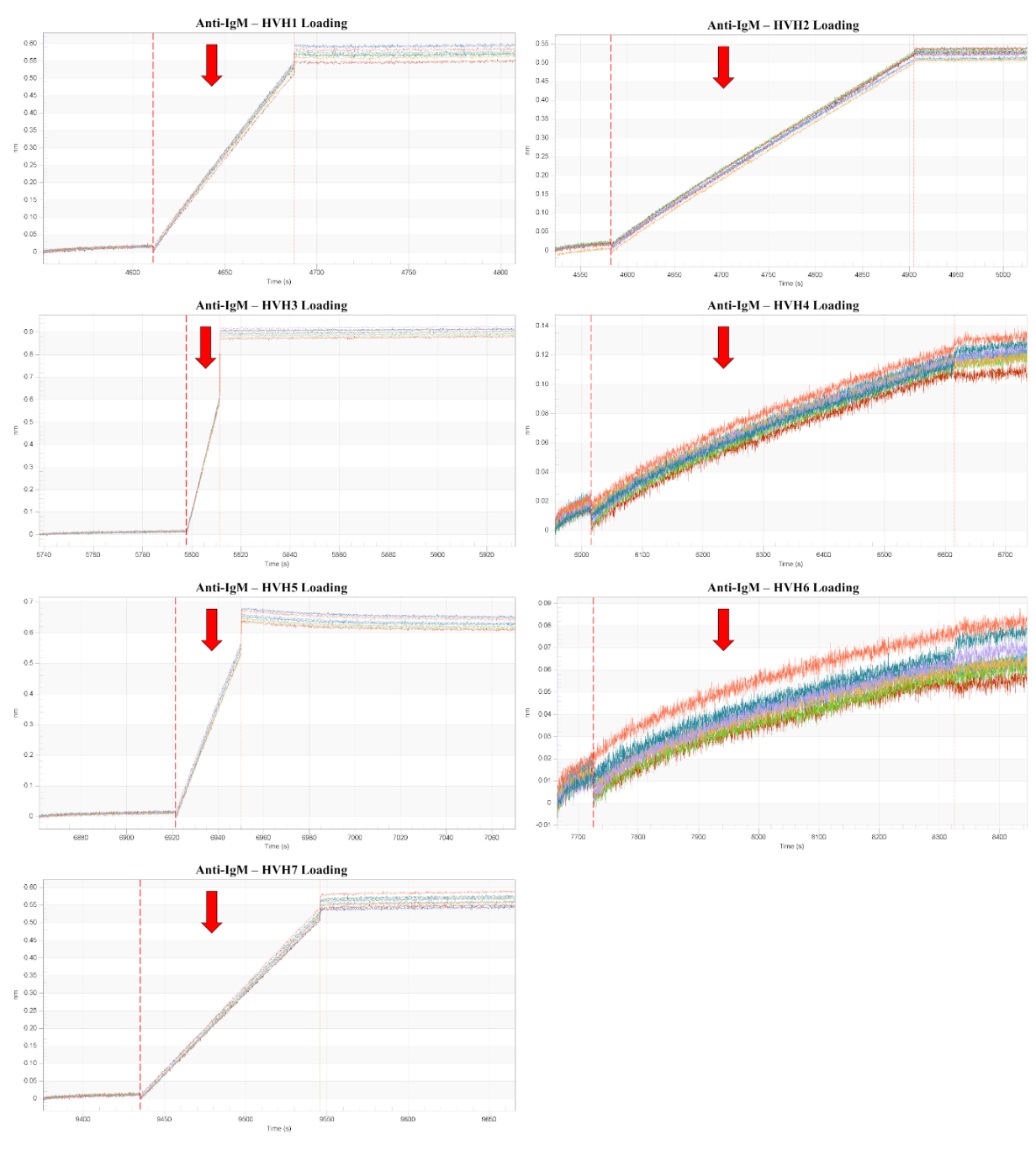

**Supplementary Figure S7. Loading graph of HVH1-7 on Anti-IgM bound on SA biosensor.**
